# A loss-of-function cysteine mutant in fibulin-3 (EFEMP1) forms aberrant extracellular disulfide-linked homodimers and alters extracellular matrix composition

**DOI:** 10.1101/2022.02.25.481983

**Authors:** DaNae R. Woodard, Steffi Daniel, Emi Nakahara, Ali Abbas, Sophia M. DiCesare, John D. Hulleman

**Affiliations:** Department of Ophthalmology, University of Texas Southwestern Medical Center, 5323 Harry Hines Blvd, Dallas, Texas, United States; Dpartment of Pharmacology, University of Texas Southwestern Medical Center, 5323 Harry Hines Blvd, Dallas, Texas, United States

**Keywords:** fibulin-3, EFEMP1, disulfide bond, extracellular matrix, marfanoid syndrome

## Abstract

Fibulin-3 (F3 or EFEMP1) is a disulfide-rich, secreted glycoprotein necessary for maintaining extracellular matrix (ECM) and connective tissue integrity. Two studies have identified distinct autosomal recessive F3 mutations in individuals with Marfan Syndrome-like phenotypes. Herein we characterized how one of these mutations, c.163T>C; p.Cys55Arg (C55R), disrupts F3 secretion, quaternary structure, and function by forming unique extracellular disulfide-linked homodimers. Dual cysteine mutants suggest that the C55R-induced disulfide species forms because of new availability of Cys70 on adjacent F3 monomers. Surprisingly, mutation of single cysteines located near C55 (i.e., C29, C42, C48, C61, C70, C159, and C171) also produced similar extracellular disulfide-linked dimers, suggesting that this is not a phenomenon isolated to the C55R mutant. To assess C55R functionality, F3 knockout (KO) retinal pigmented epithelial (RPE) were generated, followed by reintroduction of wild-type (WT) or C55R F3. F3 KO cells produced lower levels of the ECM remodeling enzyme, matrix metalloproteinase 2 and reduced formation of collagen VI ECM filaments, both of which were partially rescued by WT F3 overexpression. However, C55R F3 was unable to compensate for these same ECM-related defects. Our results highlight the unique behavior of particular cysteine mutations in F3 and uncover potential routes to restore C55R F3 loss-of-function.

## INTRODUCTION

Fibulin-3 (F3, encoded by the *EFEMP1* gene) is a secreted, disulfide rich, extracellular matrix (ECM) glycoprotein of unclear function that is broadly expressed throughout the body(Daniel et al., 2020; Zhang & Marmorstein, 2010). Increasing evidence suggests the involvement of F3 in a variety of diseases ranging from gliomas(Hu et al., 2011), to marfanoid syndrome(Bizzari et al., 2020; Driver et al., 2020), to primary open angle glaucoma(Mackay, Bennett, & Shiels, 2015), to age-related macular degeneration(Meyer et al., 2011), macular dystrophies(Hulleman, 2016; Stone et al., 1999), and general retinal dysfunction(Woodard, Nakahara, & Hulleman, 2021). Yet knowledge of the factors that influence F3 protein homeostasis is limited. Specifically, how many disease-associated mutations alter F3 folding, secretion, and degradation is largely unknown. Moreover, how these mutations influence F3 behavior will lead to a better understanding of disease mechanism and potentially rational therapies.

F3 is composed of six epidermal growth factor (EGF) domains, one of which is an atypical EGF domain containing an ∼88 residue insert(Timpl, Sasaki, Kostka, & Chu, 2003; Zhang & Marmorstein, 2010), with the remaining five domains resembling canonical calcium-binding EGF (cbEGF) domains (Fig. 1A). Structurally, each of these domains contain three disulfide bonds that form in a predictable pattern with numbering referring to the relative position of the cysteines within each domain: 1-3 [the “a” disulfide bond], 2-4 [the “b” disulfide bond], 5-6 [the “c” disulfide bond]. F3 contains 40 cysteines, at least 30 of which are engaged in disulfide bonds. Previously, we demonstrated that mutation of cysteine residues in select canonical cbEGF domains results in protein non-secretion and retention of F3 intracellularly(Hulleman, Kaushal, Balch, & Kelly, 2011; Woodard, Nakahara, et al., 2021), highlighting the importance of these residues in regulating F3 protein homeostasis. Recently, whole exome sequencing of a pair of siblings with marfanoid syndrome revealed a new homozygous cysteine missense mutation (c.163T>C; p.Cys55Arg, C55R) in F3(Bizzari et al., 2020). Patients with this homozygous mutation presented with tall stature, inguinal hernias, and advanced bone age, among other features(Bizzari et al., 2020). Due to its mode of inheritance, and because an additional study identified biallelic loss-of-function mutations in *EFEMP1* in an unrelated individual also with a Marfan-like connective tissue disorder(Driver et al., 2020), this C55R mutation likely renders F3 functionally inactive. Yet how this mutation disrupts F3 function is unknown. The C55 residue is located in the atypical EGF domain 1 (D1) which is not annotated in Uniprot to have disulfide bonds, most likely due to a lack of experimental evidence to support these assignments. Nonetheless, regardless of the *in silico* annotation, it is quite likely that the C55 residue is involved in disulfide bond formation within the atypical EGF domain and is important for F3 structural regulation. Herein, we experimentally determined how the C55R mutation alters F3 secretion, intracellular levels, and quaternary structure when produced in cultured cells. Moreover, we demonstrate that C55R F3 is unable to compensate for ECM changes driven by loss of endogenous F3, confirming that this mutation is indeed loss-of-function.

**Figure 1.**
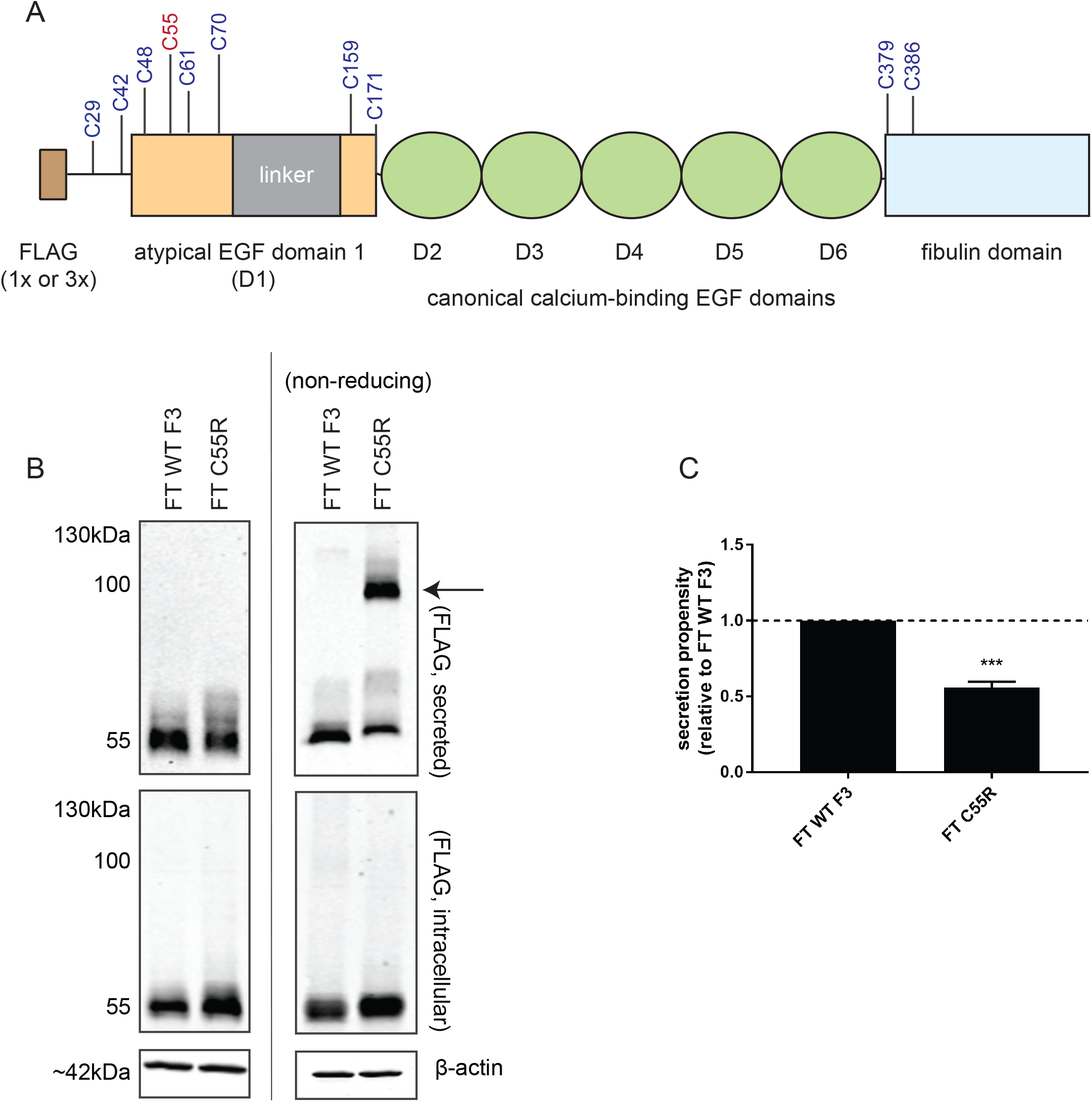
The C55R mutation in F3 results in extracellular disulfide-linked dimerization. (A) Cartoon schematic of F3 with highlighting 10 cysteines that are either in a free confirmation or are not annotated in the Uniprot database (Q12805). The pathogenic C55R mutation associated with marfanoid syndrome is highlighted in red. (B) Western blot of secreted and intracellular FLAG-tagged (FT) WT F3 and C55R variants from transfected HEK-293A cells under reducing (left) or non-reducing (right) conditions. Arrow indicates C55R dimerization. (C) Quantitation of C55R secretion propensity compared to WT F3. n = 7, mean ± SEM (*** - p<0.001, one sample t-test vs. hypothetical value of 1 [i.e., unchanged]).

## RESULTS

### Secreted C55R F3 forms an extracellular disulfide-linked dimer

Bizzari et *al*. identified a homozygous C55R mutation in the atypical EGF domain (D1, Fig. 1A) of F3 as the causative mutation resulting in marfanoid syndrome in two siblings(Bizzari et al., 2020). *A priori*, one might speculate that this mutation would cause protein non-secretion and intracellular retention, similar to what we have observed previously with engineered cysteine to alanine mutations in cbEGF domain 6 (D6) of F3 (Fig. S1A, B)(Hulleman et al., 2011; Woodard, Nakahara, et al., 2021). Surprisingly, after transfection of FLAG-tagged (FT) C55R F3 in HEK-293A cells, we found that under reducing conditions, the C55R variant was secreted at similar levels to wild-type (WT) F3 (Fig. 1B, upper left panel, 0.89 ± 0.12 vs. WT levels), and that intracellular C55R steady state levels were elevated (Fig. 1B, lower left panel 1.59 ± 0.15 vs. WT levels), resulting in a significantly reduced secretion propensity (defined as secreted/intracellular relative to WT F3 values, 0.56 ± 0.04 relative to WT, p<0.001, Fig. 1C). Yet, under non-reducing conditions, we found that in addition to a monomeric form of F3 with a slightly slower migration pattern relative to WT F3, the C55R variant also forms a secreted disulfide linked dimer at ∼110 kDa (Fig 1B, right upper panel, arrow). Under these same conditions, no detectable dimeric WT or C55R F3 was observed in the cell lysates (Fig. 1B, right middle panel). Additional experiments using untagged C55R F3 demonstrated that the media dimerization is not due to influence of the FLAG tag (Fig. S2). However, due to its versatility and cleanliness of detection (e.g., see multiple non-specific bands (*) in Fig. S2), we continued to use FLAG-tagged F3 constructs for the remainder of this work. Moreover, alkylation of conditioned media with iodoacetamide prior to boiling in non-reducing sample buffer demonstrated that this disulfide species is not forming aberrantly during sample preparation (Fig. S3). Mass spectrometry analysis of the dimer band purified by FLAG immunoprecipitation revealed that F3 was by far the most abundant protein in the 110 kDa C55R band (1.04 × 10^8^ spectral counts, Table S1), and other identified proteins (between 40 – 90 kDa) were >10-fold lower in abundance than F3 (Table S1). Overall, these data suggest that the C55R variant leads to a unique, dimerized quaternary structure primarily outside of the cell.

### The C55R mutant forms homodimers

While we have shown that C55R forms extracellular disulfide-linked dimers, it was unclear whether these dimers consisted of only C55R (homodimers) or if they also recruited WT F3 (heterodimers), both of which would be detected as F3 in the mass spectrometry results. To address this point, we performed co-transfection experiments with WT F3 and C55R containing different epitope tags (FT or 3xFT HiBiT(Dixon et al., 2016)). 3xFT HiBiT WT F3 cotransfected with FT C55R F3 was not recruited to form any detectable dimer as determined by HiBiT blotting using non-reducing conditions (Fig. 2A), indicating a lack of interaction between WT and C55R F3. However, 3xFT HiBiT C55R F3 (Fig. 2A, C) and FT C55R F3 (Fig. 2B) both formed dimeric species under the same conditions. These results are consistent with the observation that individuals heterozygous for the C55R mutation are unaffected and appear to retain sufficient activity of WT F3 to prevent marfanoid symptoms(Bizzari et al., 2020), and that this mutation does not initiate disease through a dominant negative mechanism on WT F3.

**Figure 2.**
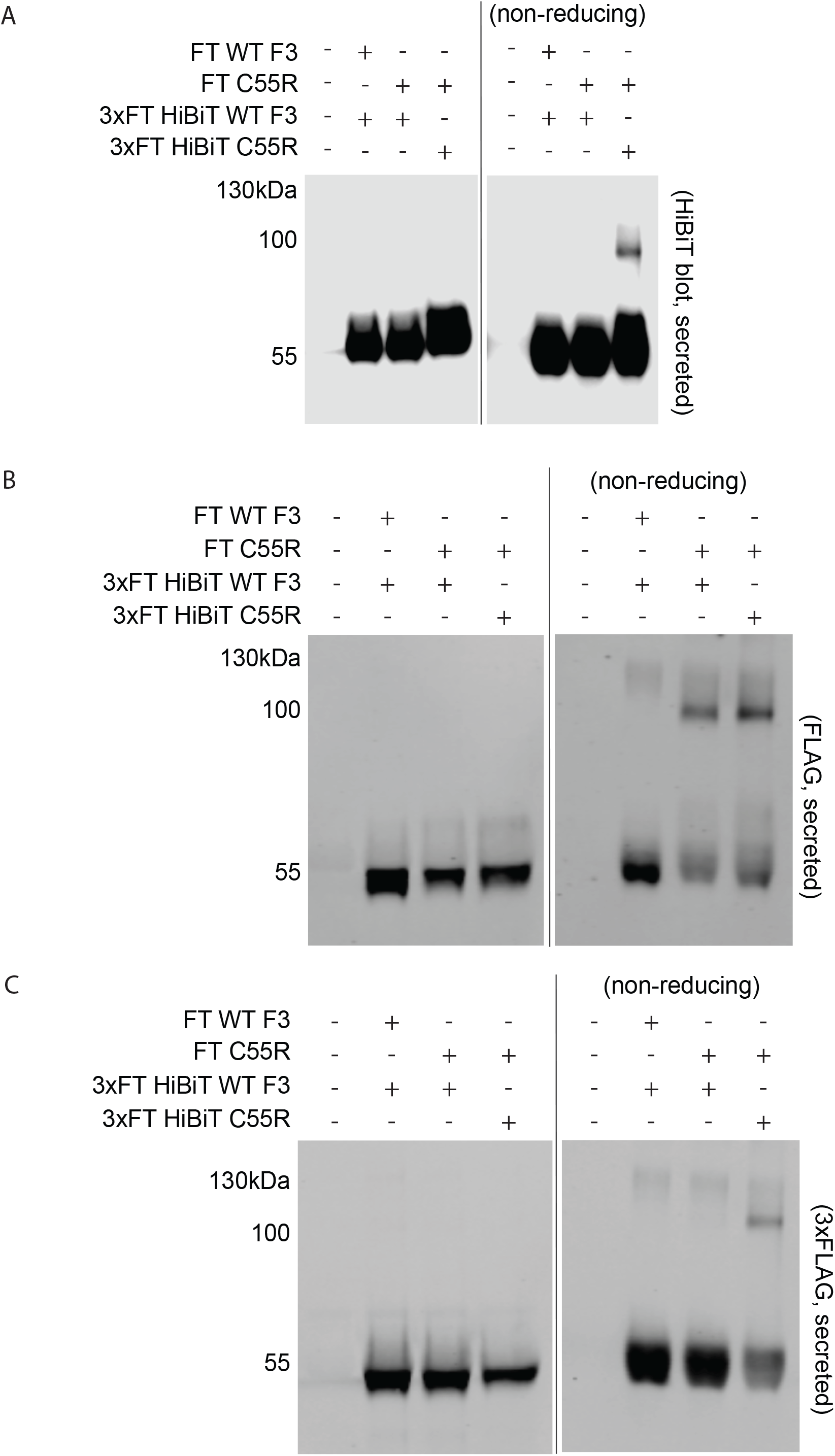
C55R forms a disulfide-linked homodimer and does not appear to sequester WT F3. (A-C) HEK-293A cells were co-transfected with constructs encoding for FT WT F3, FT C55R, 3xFT HiBiT WT F3, or 3xFT HiBiT C55R. After transfection, media was changed to serum-free and 24 h later the conditioned media was collected western blotting for (A) HiBiT (using LgBiT and NanoGlo substrate), (B) 1xFLAG (rabbit anti-FLAG, Thermo), or (C) 3xFLAG (mouse anti-FLAG M2, Sigma). Representative data of n = 3 independent experiments.

### Mutation of additional N-terminal cysteines in F3 also renders it prone to extracellular disulfide-linked dimerization

We next asked the question of whether the disulfide-linked dimer caused by the C55R mutation was unique to a mutation of this particular residue, or if it was a shared phenomenon common of other N-terminal cysteine mutants in F3. To answer this question, we generated a panel of alanine substitutions at each of the first eight cysteine residues (C29, C42, C48, C55, C61, C70, C159, and C171) spanning D1 and immediately before it (Fig. 1A). Additionally, as controls, we included alanine mutations of the last two cysteines in F3 (C379, C386), which have not been assigned disulfide bonds (Uniprot, https://www.uniprot.org/uniprot/Q12805). Each of these cysteine mutations significantly decreased the secretion propensity of F3 to a greater degree than the C55R mutation (Fig. 3A, B), suggesting that they are all important for F3 secretion and are likely engaged in transient disulfide linkages during F3 folding or in final native disulfide bonds in mature F3(Chang, Li, & Lai, 2001; Chang, Schindler, Ramseier, & Lai, 1995). Similar to the C55R variant, mutation of the remaining first five cysteines in F3 (C29, C42, C48, C61, or C70) also caused disulfide linked dimers at ∼110 kDa under non-reducing conditions, although they did so to varying degrees (Fig 3A, right upper panel, solid arrow), suggesting that formation of the dimer is not unique to only the C55R variant. Interestingly, C159A and C171A, which are located after the insertion region in D1 (Fig. 1A), formed higher molecular weight species migrating closer to ∼130 kDa (Fig. 3A, right upper panel, dashed arrow), suggesting the presence of a distinct conformation compared to the first six cysteine mutants prior to the D1 insert region. Interestingly, only a small amount of secreted monomeric F3 was found for the C379A and C386A F3 variants (Fig. 3A, right upper panel) located in the region immediately after D6 (Fig. 1A). Importantly, as with the C55R variant, the observed dimeric species for these additional cysteine mutations were also not due to aberrant disulfide bonding during sample preparation (Fig. S3). Overall, these observations suggest that quality control mechanisms regulating F3 disulfide bonding, folding and secretion appear to tolerate similar mutations vastly differently depending on their location within the F3 protein, a finding which is similar to what we have postulated when analyzing other engineered and clinically-identified mutations (Nguyen & Hulleman, 2015; Woodard, Nakahara, et al., 2021).

**Figure 3.**
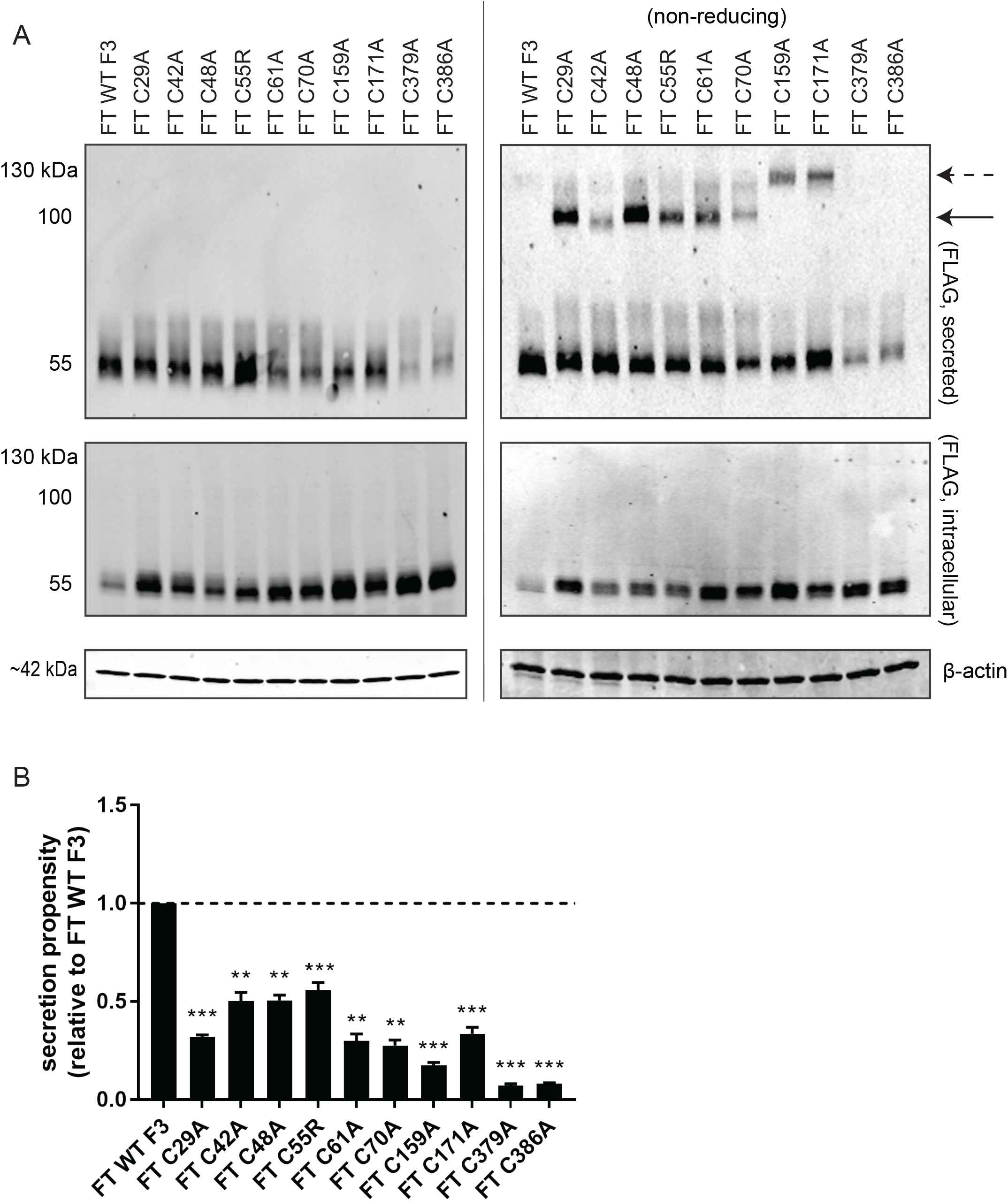
Additional cysteine mutations in F3 lead to extracellular disulfide dimer formation. (A) Western blot of cysteine variants under reducing (left panel) and non-reducing (right panel) conditions. The solid arrow indicates the 110 kDa species while the dashed arrow represents an apparent 130 kDa species. (B) Quantification of secretion propensities in (A) (reducing conditions western was quantified), n ≥ 3 independent experiments, mean ± SEM (** - p<0.01, *** - p<0.001, one sample t-test vs. hypothetical value of 1 [i.e., unchanged]).

### Dual cysteine mutations in the atypical EGF domain clarify the residues responsible for disulfide dimer formation

We speculated that the dimerization observed with the single cysteine-to-alanine mutations (Fig. 3A) in and surrounding D1 were due to new accessibility of the partner cysteine which would normally have been engaged in disulfide bonding in native F3. Thus, we hypothesized that elimination of the mutated cysteine’s disulfide bonding partner would eliminate the disulfide dimer and also shed light on the disulfide structure make up of D1. Since Uniprot defines the beginning of D1 as residue 26, disulfide bonding would be predicted to follow standard patterning for EGF domains(Arolas, Aviles, Chang, & Ventura, 2006; Chang et al., 2001; Chang et al., 1995),and the bonding order would be C29[a_n_]-C48[a_c_] (wherein the subscript indicates which residue is located closer to the N- or C-terminus), C42[b_n_]-C55[b_c_], C61[c_n_]-C70[c_c_], followed by C159[‘d’_n_]-C171[‘d’_c_] (designated as such because it seems to be an extraneous disulfide bond under this numbering, Fig. 4A). However, we hypothesized that the beginning of D1 is actually at residue 44, which is the start of exon 4 in *EFEMP1*, beginning with the canonical cbEGF leader sequence, DIDE. In fact, each of the cbEGF domains in F3 are located within their own exon (exons 5-9) and begin with the amino acids DIDE or DINE in each instance (Benchling, https://benchling.com/, ENSG00000115380). Thus, we speculate that the actual disulfide pattern is C29[a’_n_]-C42[a’_c_] (designated as such because it occurs before the a_n_ and a_c_ disulfide bond), C48[a_n_]-C61[a_c_], C55[b_n_]-C70[b_c_] followed by C159[c_n_]-C171[c_c_] (Fig. 4B). Based on this hypothesis, we predicted that in the C55R background, mutating C42 to an alanine (the supposed disulfide bonding partner according to Uniprot, Fig. 4A) would not have any effect on C55R dimer formation. Indeed, this is what we observed (Fig. 4C, arrow). However, mutation of C70 to alanine in the C55R background eliminated the ∼110 kDa dimeric species (Fig. 4C, arrow), supporting our proposed disulfide bonding pattern (Fig. 4B). Further backing of our revised disulfide bonding pattern arises from the observation that the dual C29A/C42A mutant also eliminated the ∼110 kDa species observed with the single C29A or C42A mutations (suggesting that these residues form a disulfide linkage), and that the C159A/C171A dual mutation substantially reduced the 130 kDa dimeric species typically observed in the single C159A and C171A mutants (Fig. 4C). Interestingly, in many cases, there appears to be an interplay between the 110 and 130 kDa dimeric species. For example, reduction of the 130 kDa band leads to what seems to be a small increase in the 110 kDa dimeric band and vice versa. These observations highlight the complexity of disulfide bonding in EGF domains and within the F3 protein.

**Figure 4.**
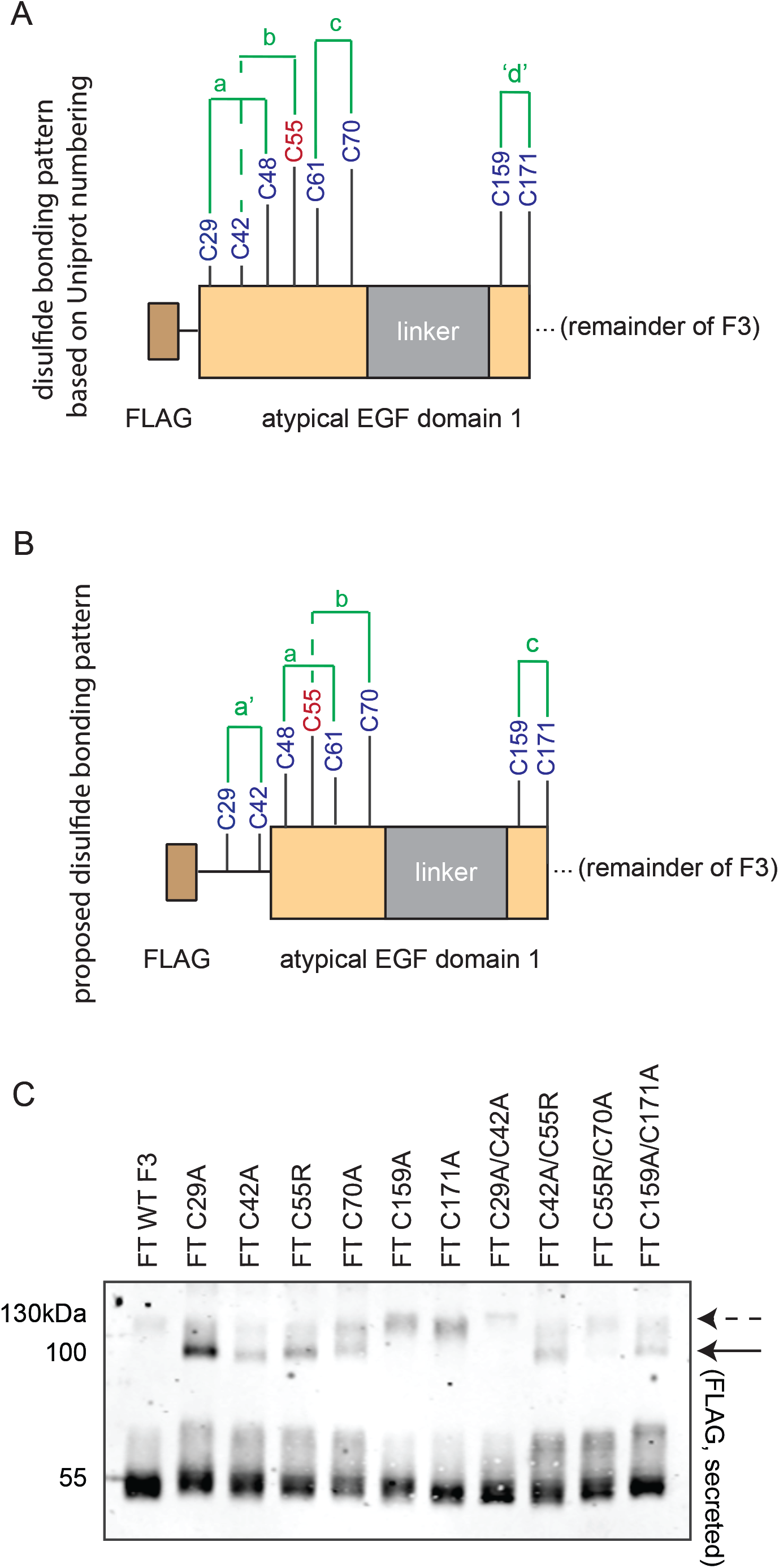
Double cysteine mutations clarify disulfide bonding patterns in the atypical EGF domain of F3. (A) The predicted disulfide bonding pattern based on where Uniprot designates the start of the atypical EGF domain (at residue 26). (B) Proposed disulfide bonding pattern based on observations that all other F3 EGF domains begin with the amino acid sequence DIDE or DINE, which makes residue 44 the beginning of this domain. (C) Non-reducing western blotting for FT F3 was performed conditioned media from transfected HEK-293A cells. The solid arrow indicates the 110 kDa species while the dashed arrow represents an apparent 130 kDa species. n = 4 independent experiments.

### F3 knockout cells demonstrate alterations in metalloproteinase activity and collagen IV assembly which are not rescuable by the C55R F3 variant

To develop a cellular background in which we could test whether C55R F3 is truly a loss-of-function mutation, we knocked out F3 in a retinal pigmented epithelial (RPE) cell line by CRISPR/Cas9 gene editing (Fig. S4A). While the physiological function of F3 is not completely clear and is most likely tissue and context-dependent, previous studies have suggested that mutations in F3 can alter matrix metalloproteineases (MMP2) and ECM formation(Fernandez-Godino, 2018; Fernandez-Godino, Bujakowska, & Pierce, 2018). Moreover, we have found that F3 KO mice have defective corneas(Daniel et al., 2020), which are largely composed of type VI collagen (ColVI)(Doane, Yang, & Birk, 1992; Zimmermann, Trueb, Winterhalter, Witmer, & Fischer, 1986). Thus, we chose to measure the activity and ECM structure of these two proteins (MMP2 and ColVI, which have also been implicated in Marfan syndrome(Ikonomidis et al., 2006; Xiong, Meisinger, Knispel, Worth, & Baxter, 2012) or myopathies(Allamand et al., 2011)), in unedited ARPE-19 cells, F3 KO cells, and F3 KO cells with overexpressed 3xFT WT F3 or 3xFT C55R F3 (Fig. S4B). F3 KO cells demonstrated a significant reduction in MMP2 activity as determined by gelatin zymography, which was partially rescued by overexpressing 3xFT WT F3 in the F3 KO background (Fig. 5A, B). However, overexpression of similar amounts of 3xFT C55R F3 did not rescue this MMP2 defect (Fig. 5A, B). Similarly, F3 KO cells showed a clear defect in ColVI ECM filament formation, which again was partly restored by 3xFT WT F3 overexpression (Fig. 5C). Expression of 3xFT C55R, on the other hand, was not able to rescue ColVI network formation (Fig. 5C). It is interesting to speculate that the asthenic and thin appearance of homozygous C55R patients(Bizzari et al., 2020) (combined with their reduced muscle strength) could be due to F3-dependent ColVI-related muscle defects(Allamand et al., 2011). The combination of these results in RPE cells strongly suggest a C55R loss-of-function mechanism.

**Figure 5.**
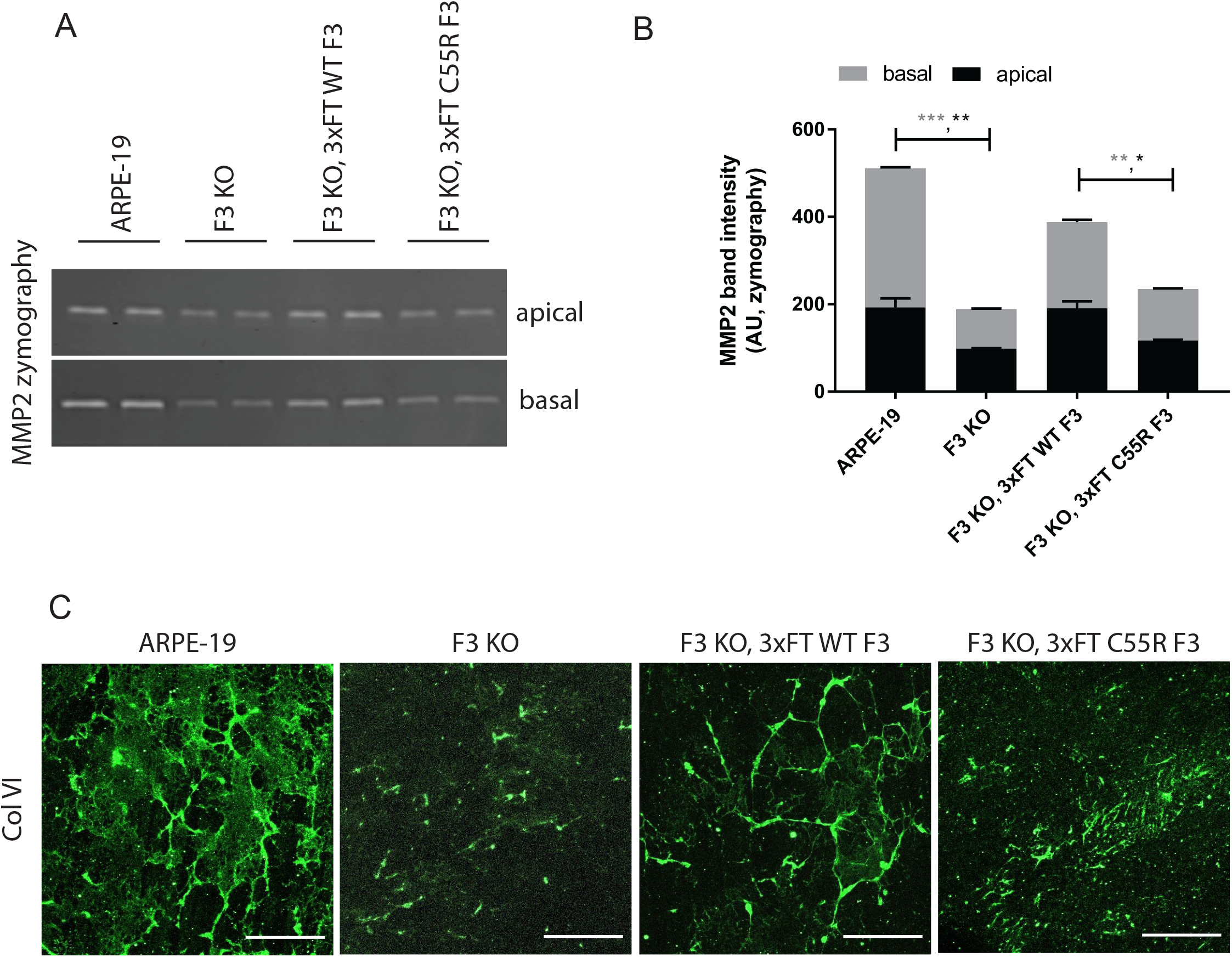
Expression of C55R F3 is unable to compensate for loss of F3, leading to changes in ECM proteins and enzymes. (A) MMP2 zymography was performed on conditioned media samples from the indicated cell lines which were polarized on transwells for 2 weeks. (B) quantification of absolute band intensity of gels shown in (A). Experiments were performed using at least biological duplicates and was performed ≥ 3 independent times. *-p<0.05, ** - p<0.01, and *** - p<0.001, two-sample unpaired t-test. (C) ColVI ECM analysis from the indicated engineered cell lines polarized on transwell inserts for 2 weeks. Transwell membranes were decellularized, fixed, stained, and imaged using confocal microscopy. Representative data of n = 2-3 independent experiments, scale bar = 50 µm.

## DISCUSSION

We report the first known instance of F3 disulfide-induced dimerization triggered by the marfanoid-syndrome-associated C55R mutation. This aberrantly formed species is detected extracellularly and appears to be a result of a C70-C70 disulfide bond on adjacent monomers. Yet, this general disulfide dimerization phenomenon does not appear to be solely unique to the C55R mutation since additional engineered cysteine-to-alanine mutations in and surrounding the atypical EGF domain region also lead to variable levels of disulfide-linked extracellular dimer. Moreover, we have demonstrated that removal of F3 from ECM-producing cells (RPE cells in this case) results in significant changes to ECM remodeling enzymes (e.g., decreased MMP2) and the morphology of prominent ECM proteins (e.g., ColVI). While overexpression of WT F3 could partially compensate for these defects, similar levels of C55R F3 overexpression did not rescue these phenotypes, reaffirming that it is a loss-of-function mutation.

The observation that multiple additional cysteine mutations (C29, C42, C48, C16, C70, C159 and C171) in the atypical EGF domain region lead to similar extracellular disulfide-linked dimers begs the question of whether additional cysteine mutations in F3 could also lead to a non-functional F3 and result in marfanoid phenotypes through similar molecular behaviors. While such mutations have not yet been identified and annotated in clinical variant databases (e.g., ClinVar, https://www.ncbi.nlm.nih.gov/clinvar/), it is quite possible that they exist in the human population, but commonly do not result in disease because of their autosomal recessive nature. Indeed, the C55R mutation would likely have not been identified if it were not for consanguinity(Bizzari et al., 2020). Our observations also raise an intriguing question of whether oxidation of certain reactive cysteine residues during normal physiology or oxidative stress could alter the structure, and therefore function of F3 and the composition/dynamics of the ECM.

In our analysis of this new C55R variant, as well as all the other cysteine mutant located near the atypical EGF domain, we were surprised to not detect any intracellular F3 disulfide bonding. While we cannot rule out the idea of intracellular disulfide formation followed by efficient secretion from the cell, the formation of extracellular disulfide bond formation appears to be a relatively unique phenomenon generally associated with coagulation (thrombus/fibrin generation)(Cho, Furie, Coughlin, & Furie, 2008; Popescu, Lupu, & Lupu, 2010). It will be interesting to more definitively test whether cysteine-mutated F3 could also be a substrate of extracellular protein disulfide isomerases.

Moving forward, our data also provide clues for potentially treating C55R-associated marfanoid syndrome. We speculate that elimination of the extracellular dimeric species may be able to restore some degree of F3 function within the ECM. To accomplish this, it could be possible to use selective, covalent cysteine nucleophiles to target and modify the two C70 residues on adjacent F3 monomers which seem to facilitate the ∼110 kDa C55R dimer formation. Such an approach, which was previously thought to be too non-specific to be useful, has gained recent momentum and promise for other proteins(Hallenbeck, Turner, Renslo, & Arkin, 2017; Resnick et al., 2019). Whether this approach would be feasible and/or beneficial in treating C55R-related marfanoid syndrome remains to be determined.

## METHODS

### Plasmid generation

Cysteine mutations were introduced into a pENTR1A FLAG (FT) WT F3 or pENTR1A 3xFT WT F3 plasmid using the Q5 Site-Directed Mutagenesis Kit (New England Biolabs, Ipswich, MA, USA). pENTR1A constructs were shuttled into the pcDNA DEST40 vector (Life Technologies, Carlsbad, CA, USA) or the pLenti CMV Puro DEST vector (a gift from Eric Campeau and Paul Kaufman, Addgene plasmid #17452) by an LR clonase II reaction (Life Technologies, Carlsbad, CA, USA) to generate the final construct used for transfection/infection. pcDNA 3xFT HiBiT F3 constructs incorporated three FLAG sequences (DYKDHDGDYKDHDIDYKDDDDK) followed by a GG<u>VSGYRLFKKIS</u> peptide (HiBiT sequence underlined) and the remainder of F3. All FLAG-containing constructs contain the FLAG sequence immediately after the signal sequence. All F3 mutations and plasmids were verified by Sanger sequencing.

### Cell culture and transfection

Human embryonic kidney cells (HEK-293A, Life Technologies, Carlsbad, CA) were cultured at 37°C with 5% CO_2_ in Dulbecco’s minimal essential medium (DMEM) supplemented with high glucose, (4.5 g/L, Corning, Corning, NY, USA), 10% fetal bovine serum (FBS, Omega Scientific, Tarzana, CA, USA), and 1% penicillin-streptomycin-glutamine (Gibco, Waltham, MA, USA). Cells were plated at a density of ∼100,000 cells/well in a 24 well plate and transfected the following day with 500 ng of midi-prepped endotoxin-free plasmid DNA (Qiagen, Germantown, MD, USA) using Lipofectamine 3000 (Life Technologies) as described previously(Woodard, Nakahara, et al., 2021; Woodard, Xing, et al., 2021). Forty-eight hours after transfection, fresh serum-free media was added. Cells were harvested and media was collected 24 h later (72 h post transfection). For cotransfection experiments, cells were plated overnight at a density of 120,000 cells/well of a 12 well plate. 500 ng of each plasmid was used combined with 1 uL of P3000 and 3 uL Lipofectamine 3000 in OptiMEM. The next day, fresh media containing 0.5% FBS was added. Media was assayed 24 h later by HiBiT blotting or standard western blotting (48 h post initial transfection). To generate stably expressing F3 cell lines, HEK-293A or ARPE-19 cells were infected with VSV-G-pseudotyped lentivirus packaged (described in more detail previously(Ramadurgum & Hulleman, 2020; Ramadurgum et al., 2020)) with the pLenti CMV Puro vector containing 3xFT WT or 3xFT C55R F3. Stable populations were selected using puromycin (1 µg/mL).

### Western and HiBiT blotting

For western blotting, cells were washed with Hanks buffered salt solution (HBSS, Sigma-Aldrich, St. Louis, MO, USA), then lysed with radioimmunoprecipitation assay (RIPA) buffer (Santa Cruz, Dallas, TX, USA) supplemented with Halt protease inhibitor (Pierce, Rockford, IL, USA) and benzonase (Millipore Sigma, St. Louis, MO, USA) for 3-5 mins at RT and spun at max speed (14,800 rpm) at 4°C for 10 min. The soluble supernatant was collected and protein concentration was quantified via bicinchoninic assay (BCA) assay (Pierce Thermo Scientific, Rockford, IL, USA). Normalized soluble supernatant or 20 μL of conditioned media was run on a 4-20% Tris-Gly SDS-PAGE gel (Life Technologies) and transferred onto a nitrocellulose membrane using an iBlot2 device (P0 program, Life Technologies). After probing for total transferred protein using Ponceau S (Sigma-Aldrich), membranes were blocked overnight in Odyssey Blocking Buffer (LICOR, Lincoln, NE, USA). Membranes were probed with rabbit anti-FLAG (1:5000; Thermo Fisher Scientific, Waltham, MA, USA, cat # PA1-984B), mouse anti-FLAG M2 (1:1000, Sigma Aldrich, St. Louis, MO, USA, cat# F1804), rabbit anti-F3 (ProSci, Poway, CA, cat# 5213), or mouse anti-β-actin (1:1000; Sigma-Aldrich, cat# A1978) for 1 h at RT followed by washes with TBS-T and an appropriate NIR-conjugated secondary antibody (LI-COR). All Western blot imaging was performed on an Odyssey CLx (LI-COR). A similar process was used for Nano-Glo HiBiT blotting (Promega, Madison, WI) wherein samples were prepared for SDS-PAGE, transferred to nitrocellulose membranes and then incubated in TBS-T for 30 min at RT. Blots were then incubated in 1x HiBiT blotting buffer supplemented with LgBiT (1:200) and Nano-Glo substrate (1:100) for 5 min at RT. Chemiluminescence was imaged using a LI-COR Fc. Images and band quantification was performed using Image Studio software (LI-COR).

### Immunoprecipitation (IP) and mass spectrometry

Serum-free media was collected from Hyperflasks (Corning) containing HEK-293A cells stably-expressing 3xFT C55R F3 and transferred to 50 ml conical tubes. A small aliquot (20 μL) of anti-FLAG M2 magnetic beads (Millipore Sigma) was added to each tube and allowed to rotate overnight at 4°C. The following day, tubes were spun at 500 rpm for 1 min. A large magnetic rack was used to capture the beads and the media was aspirated. The beads were washed three times with cold HBSS buffer supplemented with 10 mM CaCl_2_, followed by transfer to a 1.5 mL tube after the last wash. Samples were eluted with 200 μg/ml of 3xFLAG peptide (Millipore Sigma cat #F4799) in TBS supplemented with 10 mM CaCl_2_, rotated for 30 min at RT room temperature, and the supernatant was collected by placing samples in a small magnetic rack. Samples were then run on a 4-20% Tris-Gly SDS-PAGE gel (Life Technologies) followed by Coomassie Blue staining. Bands from immunoprecipitated samples were visualized using an Amersham Imager 600 (GE Healthcare), and the protein band at 110 kDa for C55R F3 was manually excised, placed in 1.5 mL Eppendorf tube, and submitted to the UT Southwestern Proteomics Core for analysis.

### F3 knockout (KO) cell generation

F3 KO cells were generated through a failed attempt to perform homology directed repair to generate point mutations in the F3 gene. Briefly, low passage ARPE-19 cells were electroporated (1400 V, 20 ms, 2 pulses, Neon Transfection System, Life Technologies) with a CRISPR/Cas9 ribonucleoprotein (RNP) containing Alt-R tracrRNA/crRNA duplex (gRNA: GACCACAAATGAATGCCGGG), Alt-R HiFi S.p. Cas9 nuclease V3, and a single stranded oligonucleotide (all from Integrated DNA Technologies, Coralville, IA). After electroporation, cells were expanded and submitted for cell sorting (UT Southwestern Flow Cytometry Core). Single colony clones were isolated, grown, and their mRNA was extracted (Aurum Total RNA Mini kit, BioRad) and analyzed for F3 expression (described previously(Woodard, Nakahara, et al., 2021)), identifying KO cells in the process. These cells were then used as the parental cells to generate the 3xFT WT F3 and 3xFT C55R F3 overexpression lines.

### Zymography

ARPE-19 cells were plated overnight at a density of 100,000 cells/well of a 12 well polyester transwell (0.4 µm pore size, 1.12 cm^2^ surface area, Corning) in full serum-containing media. The next day, the media was changed to serum-free media, and was changed every 3-4 days thereafter for 2 weeks. Prior to initiating zymography, cells were incubated in serum-free media for 72 h. Apical and basal aliquots were taken, mixed with non-reducing SDS-PAGE buffer, and loaded onto a 10% gelatin gel (9-10 µL for apical, 18-20 µL for basal, Novex, Life Technologies). Samples were electrophoresed for ∼90 min at 125-140V. The gel was then incubated in renaturing buffer (30 min at RT, Novex), followed by incubation in developing buffer (30 min at RT, G-Biosciences, St. Louis, MO), and again in fresh developing buffer overnight at 37°C with gentle shaking. Gels were stained with Coomassie R-250 for 1 h at RT, followed by destaining to reveal digested gelatin corresponding to MMP2 activity. Gels were imaged on an Odyssey CLx (LI-COR) using the 700 nm channel and quantified using Image Studio (LI-COR).

### Extracellular matrix (ECM) analysis

Cells were seeded on transwells and treated as described above in the zymography section. After 2 weeks, transwells were washed 3x with DPBS (Gibco) and decellularized by incubation with 0.5% Triton-X 100 and 20 mM ammonium hydroxide for 5 min at 37°C with gentle shaking(Fernandez-Godino et al., 2018). Samples were washed 1x in DPBS followed by fixation in 4% PFA for 30 min at RT. Transwells were then washed 2x in DPBS and blocked for 1 h at RT in 10% goat serum, 0.1% BSA, 0.1% Triton-X dissolved in PBS. Primary antibody (rabbit anti-ColVI, 1:200, Abcam #ab6588) was incubated at 4°C overnight with gentle rocking. Membranes were washed 2x followed by secondary antibody (anti-rabbit AlexaFluor 488 or 594, 1:1000, Life Technologies), for 2 h at RT. After two additional DPBS washes, DAPI was added (to confirm decellularization), incubated for 5-10 min at RT and then transwells were washed again one final time before excision and mounting on glass slides with a drop of ProLong Diamond Antifade (Life Technologies). Images were acquired at either 63x or 25x magnification using a Leica TCS SP8 confocal microscope (Leica Microsystems) and analyzed using ImageJ (NIH). Images presented were acquired by z-scanning (∼15 steps, 0.37 µm each) at 63x magnification.

### Statistical analysis

To determine statistical significance, samples were compared using either a one-sample t-test using Excel against a hypothetical value of 1 (i.e., unchanged compared to the control), or a two-sample unpaired t-test. Significance was set at *p - <0.05, **p - <0.01, and *** -p<0.001.

## FIGURE LEGENDS

**Table S1. List of proteins identified by mass spectrometry in the C55R dimer band**.

**Figure S1. Cysteine to alanine mutations in domain 6 of F3 result in protein non-secretion and intracellular retention**. (A) HEK-293A cells were transfected with either FT WT F3 or a FT C338A F3 construct that eliminates a known disulfide bond within domain 6 of F3 and analyzed by reducing or non-reducing western blotting. (B) Quantification of FT C338A secretion propensity. n = 3 independent experiments, mean ± SEM (*** - p<0.001, one sample t-test vs. hypothetical value of 1 [i.e., unchanged]).

**Figure S2. Untagged C55R F3 also forms extracellular dimers**. HEK-293A cells were transfected with either untagged WT F3 or untagged C55R F3. The next day, media was switched to serum-free and cells were incubated for 24 h. Conditioned media was collected and concentrated from 1.5 mL to ∼75 µL (3,000 MWCO, Millipore) and run on a non-reducing western blot. Three independent experiments are shown on the same blot. * = non-specific staining.

**Figure S3. Alkylation of conditioned media prior to boiling in non-reducing SDS-PAGE buffer does not prevent dimer formation**. Conditioned media from cells transfected with the indicated FT F3 constructs were treated with iodoacetamide (or left untreated) prior to boiling the sample in non-reducing SDS-PAGE buffer and western blotting. n = 3 independent experiments.

**Figure S4. qPCR and western blot characterization of newly engineered ARPE-19 cell lines**. (A) Messenger RNA was extracted from ARPE-19 cells, ARPE-19 F3 KO cells, or F3 KO cells stably overexpressing 3xFT WT F3 or 3xFT C55R F3 and analyzed for F3 expression. (B) Western blot of apical or basal media aliquots (serum-free) taken from transwell-polarized ARPE-19 cells used in MMP2 zymography and ColVI ECM experiments. Representative data of n = 2 independent experiments.

## ACKNOWLEDGEMENTS

JDH is supported by an endowment from the Roger and Dorothy Hirl Research Fund, and R01 EY027785. DRW was supported by an NIH Diversity Supplement. AA was supported by an NIH T35 Training Grant (T35 EY026510). Additional support was provided by a National Eye Institute Visual Science Core Grant (P30 EY030413, to the UT Southwestern Department of Ophthalmology).

## DATA AVAILABILITY STATEMENT

Data provided in this publication is available upon request to corresponding author.

