## Supplementary figures and images for "A loss-of-function cysteine mutant in fibulin-3 (EFEMP1) forms aberrant extracellular disulfide-linked homodimers and alters extracellular matrix composition"

### Fig. S1

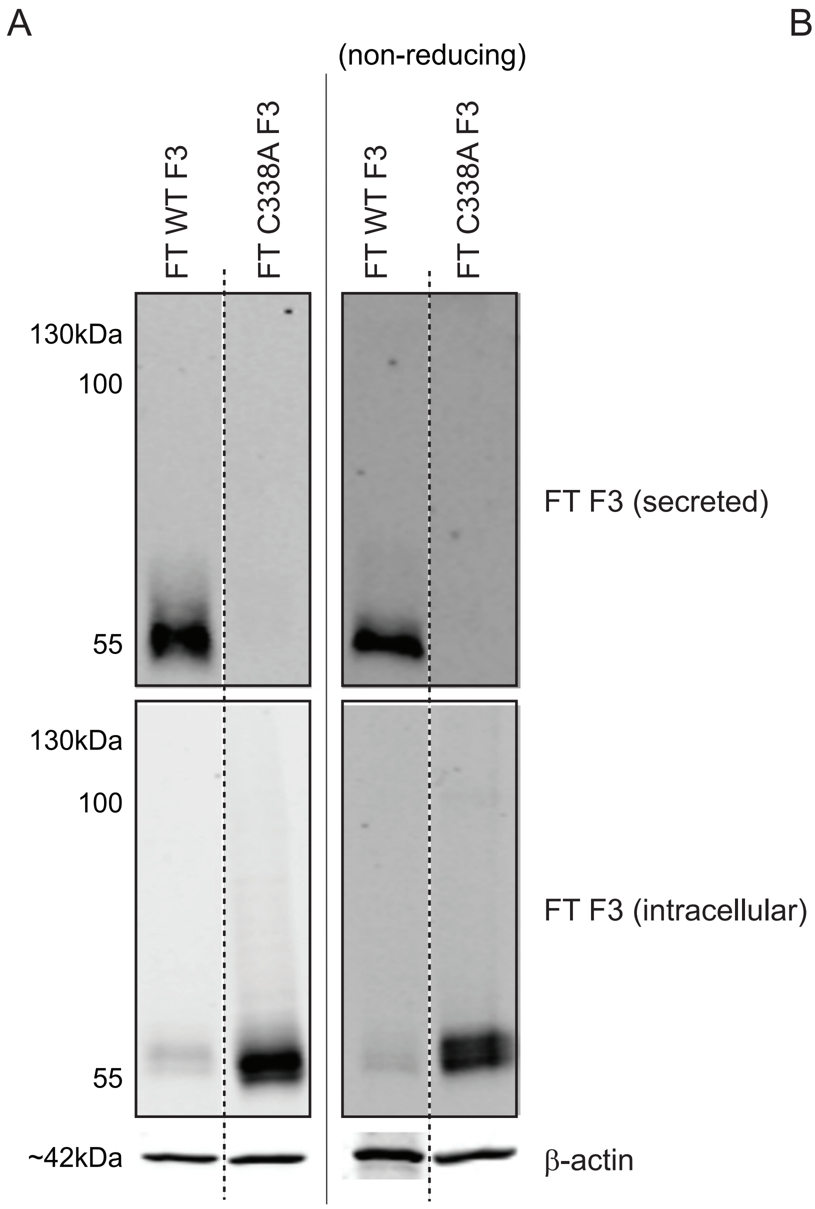

### Fig. S2

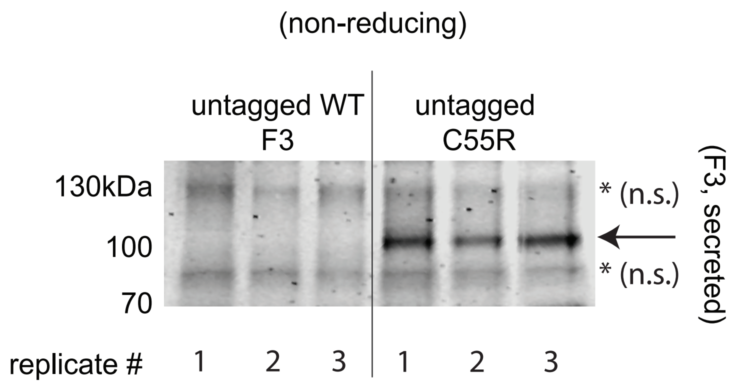

### Fig. S3

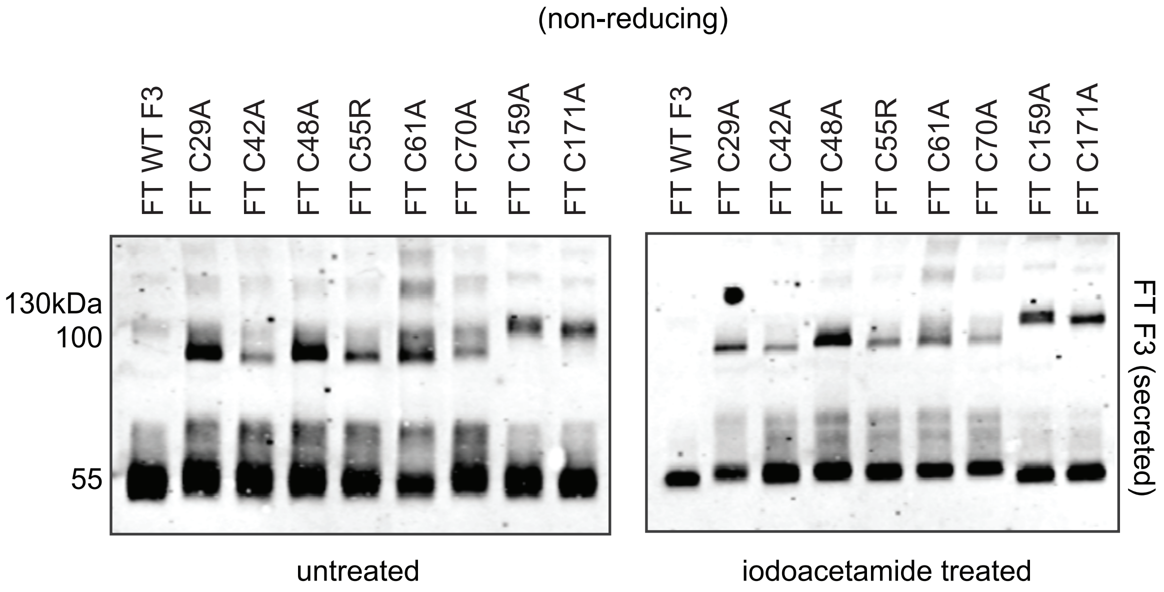

### Fig. S4

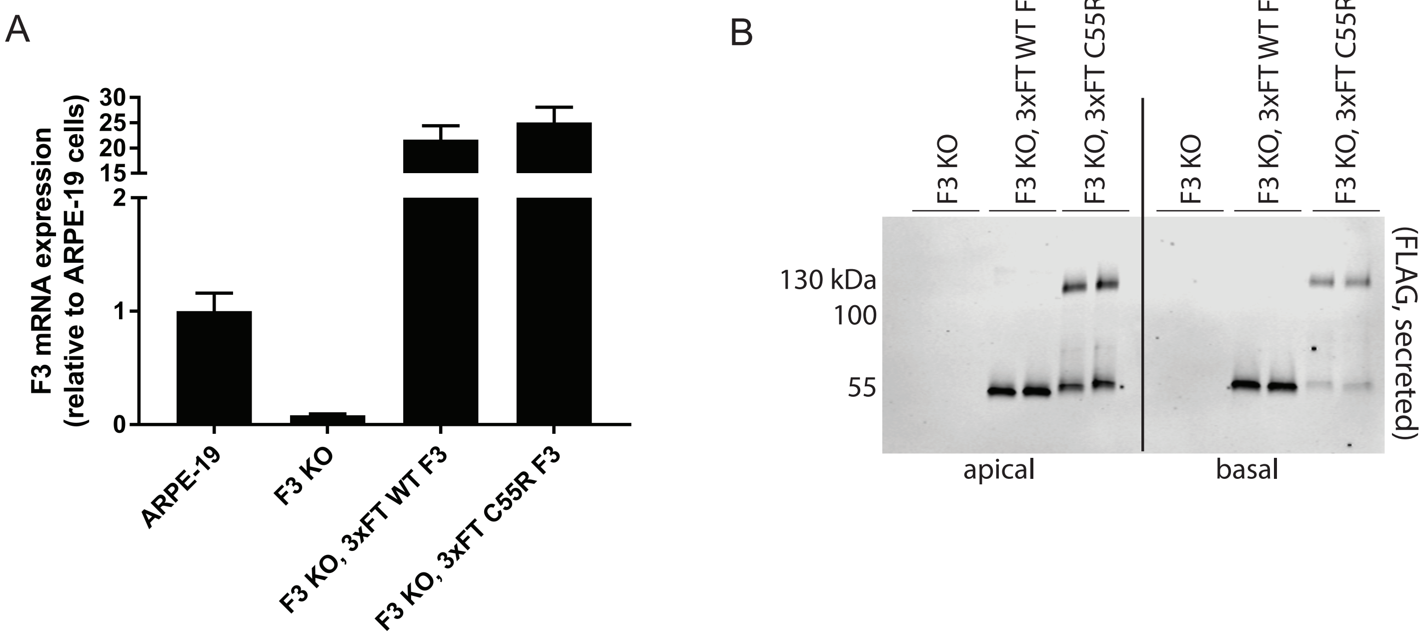
