## Supplementary material for "A loss-of-function cysteine mutant in fibulin-3 (EFEMP1) forms aberrant extracellular disulfide-linked homodimers and alters extracellular matrix composition": Table S1

| Accession | Description | Coverage [%] | # PSMs | MW [kDa] | Abundance |
|---|---|---|---|---|---|
| Q12805 | EGF-containing fibulin-like extracellular matrix protein 1 OS=Homo sapiens OX=9606 GN=EFEMP1 PE=1 SV=2 | 52 | 265 | 54.6 | 1.04E+08 |
| P02768 | Serum albumin OS=Homo sapiens OX=9606 GN=ALB PE=1 SV=2 | 8 | 47 | 69.3 | 9.48E+06 |
| P63261 | Actin, cytoplasmic 2 OS=Homo sapiens OX=9606 GN=ACTG1 PE=1 SV=1 | 36 | 22 | 41.8 | 1.53E+06 |
| F5H5D3 | Tubulin alpha chain OS=Homo sapiens OX=9606 GN=TUBA1C PE=1 SV=1 | 12 | 4 | 57.7 | 9.08E+05 |
| P13473 | Lysosome-associated membrane glycoprotein 2 OS=Homo sapiens OX=9606 GN=LAMP2 PE=1 SV=2 | 2 | 2 | 44.9 | 9.01E+05 |
| A0A1X7SBS1 | Heterogeneous nuclear ribonucleoprotein U OS=Homo sapiens OX=9606 GN=HNRNPU PE=1 SV=1 | 4 | 10 | 81.7 | 8.04E+05 |
| F8VX11 | Eukaryotic translation initiation factor 4B (Fragment) OS=Homo sapiens OX=9606 GN=EIF4B PE=1 SV=1 | 7 | 8 | 41.4 | 7.48E+05 |
| A0A087WVQ9 | Elongation factor 1-alpha 1 OS=Homo sapiens OX=9606 GN=EEF1A1 PE=1 SV=1 | 8 | 5 | 47.9 | 5.63E+05 |
| P49257 | Protein ERGIC-53 OS=Homo sapiens OX=9606 GN=LMAN1 PE=1 SV=2 | 4 | 2 | 57.5 | 2.64E+05 |
| F5GZS6 | 4F2 cell-surface antigen heavy chain OS=Homo sapiens OX=9606 GN=SLC3A2 PE=1 SV=1 | 2 | 1 | 64.8 | 1.99E+05 |
| A0A0G2JIW1 | Heat shock 70 kDa protein 1B OS=Homo sapiens OX=9606 GN=HSPA1B PE=1 SV=1 | 5 | 4 | 70.1 | 1.59E+05 |
| P60842 | Eukaryotic initiation factor 4A-I OS=Homo sapiens OX=9606 GN=EIF4A1 PE=1 SV=1 | 2 | 1 | 46.1 | 1.37E+05 |
| A0A087WW43 | Inter-alpha-trypsin inhibitor heavy chain H3 OS=Homo sapiens OX=9606 GN=ITIH3 PE=1 SV=1 | 1 | 3 | 75 | 7.43E+04 |
| B0QYK0 | RNA-binding protein EWS OS=Homo sapiens OX=9606 GN=EWSR1 PE=1 SV=1 | 9 | 7 | 64.9 | 6.95E+04 |
| P10809 | 60 kDa heat shock protein, mitochondrial OS=Homo sapiens OX=9606 GN=HSPD1 PE=1 SV=2 | 3 | 2 | 61 | 6.33E+04 |
| P12277 | Creatine kinase B-type OS=Homo sapiens OX=9606 GN=CKB PE=1 SV=1 | 4 | 1 | 42.6 | 6.21E+04 |
| O43242 | 26S proteasome non-ATPase regulatory subunit 3 OS=Homo sapiens OX=9606 GN=PSMD3 PE=1 SV=2 | 3 | 1 | 60.9 | 5.41E+04 |
| P11279 | Lysosome-associated membrane glycoprotein 1 OS=Homo sapiens OX=9606 GN=LAMP1 PE=1 SV=3 | 4 | 1 | 44.9 | 2.81E+04 |
| P35637 | RNA-binding protein FUS OS=Homo sapiens OX=9606 GN=FUS PE=1 SV=1 | 3 | 4 | 53.4 | 1.81E+04 |
